# Identification of an Outbreak of Bivalve Transmissible Neoplasia in Soft-Shell Clams (*Mya arenaria*) in the Puget Sound Using Hemolymph and eDNA Surveys

**DOI:** 10.1101/2024.12.03.626659

**Authors:** Sydney A. Weinandt, Zachary J. Child, Dorothy Lartey, Angel Santos, Holden Maxfield, Jordana K. Sevigny, Fiona E. S. Garrett, Peter D. Smith, Rachael M. Giersch, Samuel F. M. Hart, Franchesca Perez, Lucas Rabins, Samuel Kaiser, Anna Boyar, Jan Newton, Jesse Kerr, James L. Dimond, Michael J. Metzger

## Abstract

Bivalve transmissible neoplasia (BTN) is one of three known types of naturally transmissible cancer— cancers in which the whole cancer cells move from individual to individual, spreading through natural populations. BTN is a lethal leukemia-like cancer that has been observed throughout soft-shell clam (*Mya arenaria*) populations on the east coast of North America, with two distinct sublineages circulating at low enzootic levels in New England, USA, and Prince Edward Island, Canada. Major cancer outbreaks likely due to *Mya arenaria* BTN (MarBTN) were reported in 1980s and the 2000s and the disease has been observed since the 1970s, but it has not been observed in populations of this clam species on the US west coast. In 2022, we collected soft-shell clams from several sites in Puget Sound, Washington, USA, and unexpectedly found high prevalence of BTN in two sites (Triangle Cove on Camano Island and near Stanwood in South Skagit Bay). Prevalence of BTN increased in subsequent years, reaching >75% in both sites in 2024, while it was not observed in other sites, suggesting the early stages of a severe disease outbreak following recent introduction. We observed that these cancer cells contain several somatic transposing insertion sites found only in the USA-sublineage of MarBTN, showing that it likely was recently transplanted from New England to this location. We then developed a sensitive environmental DNA (eDNA) assay, using qPCR to target somatic mutations in the MarBTN mitogenome, and showed that MarBTN can be detected in seawater at Triangle Cove, as well as several kilometers outside of the cove. We then used this assay to survey 50 sites throughout Puget Sound, confirming that the disease can be detected at high levels at Triangle Cove and South Skagit Bay, and showing that it extends beyond these known sites. However, while normal soft-shell clam mtDNA was widely detected, MarBTN was undetectable throughout most of Puget Sound and currently remains limited to the South Skagit Bay area and north Port Susan. These results identify a previously unknown severe outbreak of a transmissible cancer due to long-distance transplantation of disease from another ocean, and they demonstrate the utility of eDNA methods to track the spread of BTN through the environment.

## INTRODUCTION

Most known cancers arise within an animal and remain within the same individual until the end of their life; however, there are a few rare types of horizontally transmitted cancers in which the cancer cells from one individual are able to directly infect and engraft in other animals. Transmissible cancers were first found in dogs as canine transmissible venereal tumor disease (CTVT) (1, 2) and Tasmanian devils as devil facial tumor disease (DFTD) (3). These tumors are transmitted between individuals through physical contact, and in both cases exhibit genotypes that do not match their hosts but rather show a single clonal lineage in dogs, and two independent clonal lineages in devils (4). In many bivalve species, a leukemia-like disseminated neoplasia (DN) has been reported to be a bivalve transmissible neoplasia (BTN) (5-7), with at least 10 independent lineages observed affecting at least 10 different species (8-13).

BTN is characterized by the large number of cancer cells in the hemolymph, the circulatory fluid of the bivalve, followed by dissemination into the solid tissues of the animal as the disease progresses (14, 15). These cancer cells are rounded, non-adherent, and usually polyploid, in contrast to healthy bivalve hemocytes. BTN cells are capable of surviving in artificial sea water for weeks, and detection of BTN-specific DNA in tank water of diseased clams in aquaria has shown that cancer cells are released by diseased animals, indicating that the method of transmission is likely through the water column (16-18).

DN was first reported in the 1970s in soft-shell clams (*Mya arenaria*) on the east coast of North America (19, 20). BTN is the likely cause of the severe outbreaks of DN that were reported in these populations that reached up to 90% prevalence with severe population losses in New England in the 1980s and in Prince Edward Island, Canada in the 2000s (21, 22). All samples of DN from soft-shell clams on the east coast that have been analyzed with genetic markers come from a single lineage of cancer (termed MarBTN), with two discrete sublineages in clams from USA and from Prince Edward Island (PEI) that appear to have diverged ∼200 years ago (23). Soft-shell clams are native to the east coast of North America, and to date, BTN has only been reported in these populations, despite the presence of soft-shell clams in Washington State and Europe. The current population in Washington State is not native, and was likely intentionally introduced from the Atlantic populations of clams in the 1870s (24). These circumstances appeared to provide the opportunity to study an entirely naive population of soft-shell clams on the west coast to compare with the infected soft-shell clams on the east coast. However, when we sampled 47 animals from Triangle Cove, WA in 2022 (a location with anecdotal reports of recent die-offs), we unexpectedly found clams with a high prevalence of BTN. These findings prompted us to investigate the presence of BTN in Puget Sound, WA.

Environmental DNA (eDNA) is increasingly used in marine and aquatic conservation efforts, especially in detecting invasive species and pathogens (25-30). eDNA sampling involves filtering water to capture the DNA present in the water column, which is then extracted and amplified for analysis to identify species present, or in this case cancerous cells. When used together with highly sensitive assays such as qPCR or ddPCR, eDNA samples can be used to detect trace amounts of target taxa DNA (31). While not eliminating the need for traditional sampling methods, eDNA sampling is an effective supplementary tool in monitoring efforts (25). Previous studies have demonstrated the ability of eDNA analysis to detect a variety of marine pathogens and parasites, such as MSX, perkinsosis (32), *Vibrio spp. (33)*, and abalone withering syndrome (29). One study has investigated the use of eDNA for the detection of transmissible cancers, showing clear detection in tank water of clams in aquaria (17). Here we build on those eDNA protocols and develop a more sensitive eDNA extraction and amplification method that enables sensitive detection in wild samples.

The goal of this study was twofold. First, we report the identification of MarBTN on the western coast of the United States, and we test whether it represents spread of MarBTN from the eastern coast of the United States or a new lineage. Secondly, we used both animal collection and eDNA surveys to observe the disease in the environment throughout the Puget Sound, to determine the extent of the spread of this outbreak and develop novel tools for disease monitoring.

## MATERIALS AND METHODS

### Clam collection and sampling

Adult soft-shell clams were collected and sampled from five locations in Puget Sound, Washington (Triangle Cove, Crandall Spit, near Stanwood, Similk Bay, and Sequim Bay). From each site, between 3 to 66 soft-shell clams were collected once each year from 2022 to 2024 (Table S1). Animals were collected from the intertidal zone during low tides with a shovel for all sites except clams near Stanwood, which were collected by a commercial supplier. All animals were measured, and 0.5-1 ml of hemolymph was taken from the pericardial region using a 0.5 in 26-gauge needle fitted on a 3 mL syringe (17). Approximately 20-50 μl of hemolymph was placed in a 96-well plate and incubated at 4°C for approximately 1 h before screening for the morphologically distinct cancer cells on an inverted phase-contrast microscope (Fig. 1). Hemolymph was spun down in a centrifuge chilled to 4°C at 1,000 × g to pellet the cells. The supernatant was removed, and the samples were kept at –80°C until DNA extraction. Hemolymph DNA was extracted using the Monarch Genomic DNA Purification kit (New England BioLabs).

**Figure 1.**
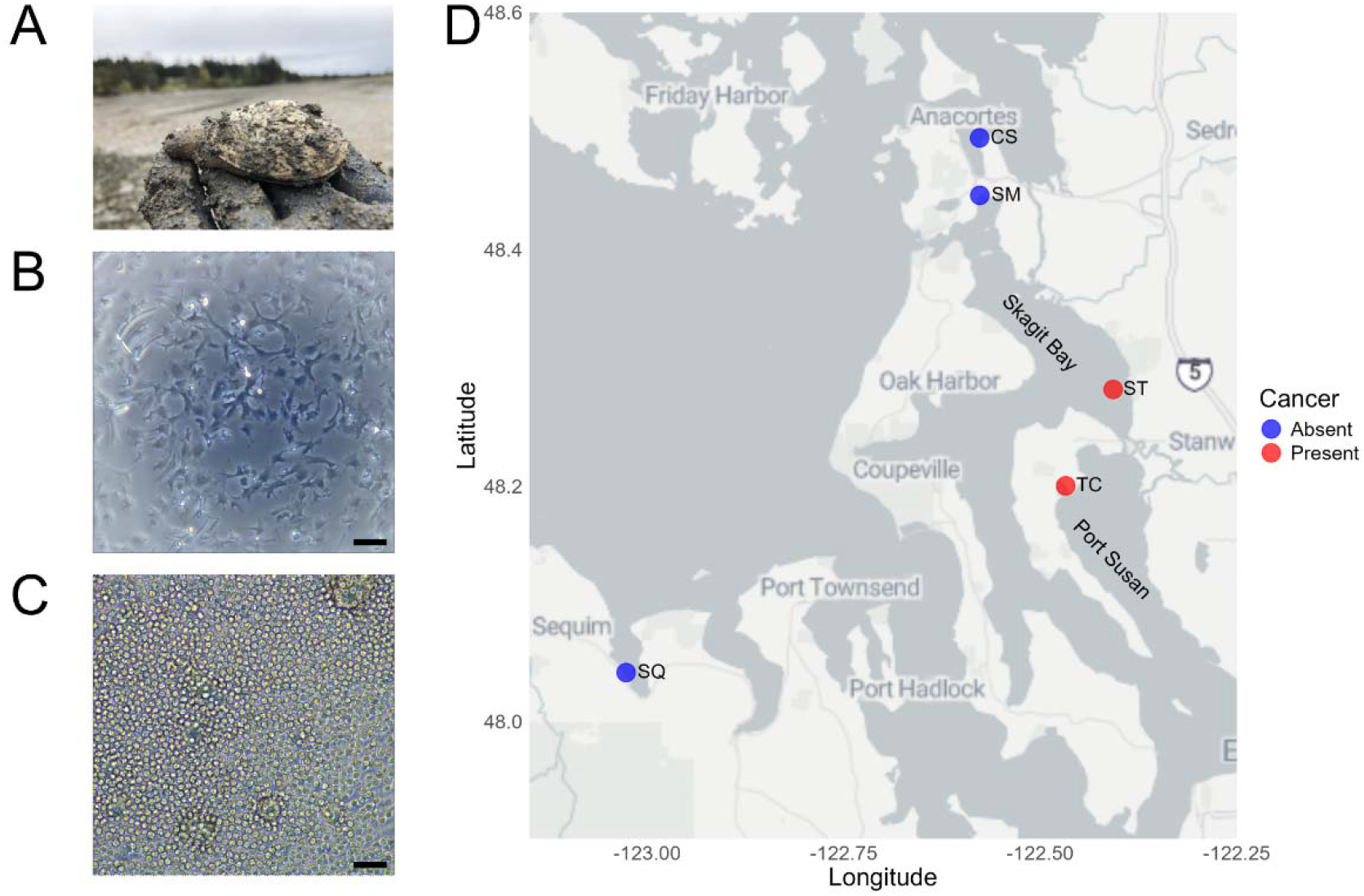
Detection of Bivalve Transmissible Neoplasia (BTN) in soft-shell clams (*Mya arenaria*) in Puget Sound, Washington State, USA. (***A***) Representative soft-shell clams (*Mya arenaria*) from Triangle Cove, Washington, USA, collected in 2022 are shown. (***B***) Representative healthy hemocytes from a clam from Crandall Spit, WA (WACS-B2) and (***C***) Bivalve Transmissible Neoplasia from a clam from Triangle Cove, WA (WATC-D5) show very different morphology. Scale bar is 50 μm. (***D***) Map shows the locations of clam collections with colors denoting whether BTN is present (red) or absent (blue). Samples were collected from 2022-2024. BTN was detected in clams at Triangle Cove (TC) and in Stanwood (ST) 2022-2024. BTN was never detected in clams from other locations at any collection 2022-2024. Specific prevalence of BTN at each collection is shown in Table 1. Map was generated using R packages ggplot, ggmap, and dplyr. Map style wa generated by StadiaMaps, using map type “alidade_smooth.”

### MarBTN qPCR diagnostic targeting a nuclear marker

Allele specific qPCR assays were conducted to quantify the neoplastic and genomic DNA in the hemolymph. One pair of primers was used to target a nuclear locus, with the cancer-specific pair targeting a somatic insertion of the LTR-retrotransposon *Steamer* near the *N1N2* gene and the “universal” clam pair targeting a conserved region of the *N1N2* ORF nearby (17). The plasmid pCR-SteamerLTR-N1N2, which contains a single copy of the locus from a cancer cell that has the targets for both cancer and control primers, was used for the standard curve. The plasmid was linearized using 0.5 μL of NotI-HF (New England Biolabs) for 2 h at 37°C in a 10 μL reaction, and heat-inactivated at 65°C for 20 min. DNA concentration was determined using Qubit, copy number per μL was calculated based on total plasmid size, and stocks were diluted to 1 × 109 copies/μL with NEB Elution Buffer. Reactions were run using PowerUp SYBR green 2x master mix (ThermoFisher Scientific) on a StepOnePlus real-time PCR cycler with a holding stage at 95°C for 2 min, two step cycling at 95°C for 15 s and 60°C for 30 s for 40 cycles, followed by the melt curve stage. All samples were run in triplicate and values were averaged. Samples were considered positive if amplification was detected in all three wells. Using these values, we used the ratio (R) of the total copies of the cancer-specific allele and the total copies of *N1N2* present to estimate the fraction of MarBTN cells in the clam hemolymph. The estimated fraction is R/(1-R) to adjust for the difference in ploidy of the diploid normal cells to the polyploid MarBTN cancer cells, which are tetraploid at this locus in the USA sublineage based on genomic analysis (23).

### PCR

To compare the genotype of MarBTN on the west coast of the US to those from the east coast of the US and Canada, we used 17 different primers that target specific insertion sites of the LTR-retrotransposon *Steamer* paired with a single forward primer that matches the end of the *Steamer* LTR (ClamLTR-F2). Previously reported primers were identified through inverse-PCR cloning strategies using genomic DNA of MarBTN samples (6, 34), and new primers were designed based on analysis of *Steamer* insertion sites from whole genome sequencing analysis (23). Primers are listed in Table S2. For each 25 μl reaction, 2.5 μl of DNA was used. Thermocycler conditions were: 95°C for 5 min, 35 cycles of 95°C for 30 s, 50°C for 30 s, and 72°C for 30 s, followed by 72°C for 5 min. The amplified DNA products were analyzed using 2% agarose gel electrophoresis with 10 μl of product per reaction. Additionally, the mitochondrial cytochrome oxidase b sequence was amplified using previously reported primers (35).

### Field seawater eDNA collection, filtration, and DNA extraction

*Shore sampling:* For collection, filtration, and extraction of eDNA from shore, seawater was collected by hand at low tide from each collection site into 3 × 500 mL separate autoclaved polypropylene laboratory bottles. Samples were held in a cooler on ice until return to the lab 2-4 h later, after which they were frozen at –30°C until filtration using a 0.45 μm cellulose nitrate filter and an autoclaved, bench-top vacuum filtration apparatus. Each 500 ml biological replicate from each collection was filtered and extracted separately. The unused outer ring of each filter was trimmed off, and the filter was cut into 6 pieces to facilitate contact with the beads during extraction. Filters were stored at –80°C until DNA extraction with DNeasy PowerSoil Pro Kits (Qiagen).

*Sampling by ship:* Water samples were collected from several cruises conducted in 2024: a collection throughout the Skagit Bay, a collection in Tulalip Bay, and a larger survey of Puget Sound conducted on the R/V Carson with the Washington Ocean Acidification Center (WOAC). Due to conditions on a ship and protocols of the WOAC, modifications to the eDNA extraction are as follows:

For Skagit Bay and the detailed collections around Triangle Cove, seawater samples were collected from one meter off the seafloor with a handheld Niskin bottle. Proper depth was determined by attaching a one-meter line with a weight on the bottom end to the Niskin bottle. Seawater was collected from each site into 3 × 500 mL separate autoclaved polypropylene laboratory bottles in Skagit Bay. For the analysis of local spread around Triangle Cove, 1 × 500 mL bottle was collected at each site at two different times in the same day. Collection bottles and the Niskin bottle were bleached and rinsed in between sampling stations. Each sample was immediately filtered through a 0.45 μm cellulose nitrate filter using Single Use Analytical Filter Funnels (Nalgene). Filters were then placed into sterile coin envelopes, and subsequently sealed in a mylar ziplock bag with a silica gel desiccant packet for short-term preservation. Upon return to the laboratory, filters were held at –80°C until extraction with DNeasy PowerSoil Pro Kits (Qiagen).

For Tulalip Bay, 3 × 500 mL seawater samples were collected from one meter off the seafloor with a handheld Niskin bottle as for Skagit Bay collections. The Niskin bottle was thoroughly rinsed with seawater upon arrival at the following sampling station. As with samples collected from shore, water samples were held in a cooler on ice until return to the lab 2-4 h later, after which they were frozen at – 30°C until filtration through a 0.45 μm cellulose nitrate with an autoclaved, bench-top vacuum filtration apparatus, and filters were held at –80°C until extraction with DNeasy PowerSoil Pro Kits (Qiagen), as described above.

For WOAC cruise on R/V Carson: Seawater samples were collected by a Niskin rosette, attached to the ship’s CTD. Three Niskin bottles were used for each site, with one programmed to close at the bottom of the cast, one in the middle, and one at the surface of the water. For each bottle, exact closure depth was recorded. 500 mL of seawater was collected from each Niskin and transferred into a 500 mL autoclaved polypropylene laboratory bottle. Each sample was immediately filtered through a 0.45 μm cellulose nitrate filter using Single Use Analytical Filter Funnels (Nalgene). Filters were then placed into 900 μL of Longmire’s solution, a lysis buffer which preserved samples until DNA extraction (36, 37), and eDNA was later extracted with a phenol:chloroform:isoamyl alcohol protocol (modified from (38)). Modifications to the published protocol were as follows: filter membranes were incubated at 56°C for 2 hours before the addition of 900 μL of phenol:chloroform:isoamyl alcohol, after which samples were subsequently both shaken vigorously and vortexed for 10 s. Glycogen blue was added immediately after transfer of the aqueous layer of the final chloroform:isoamyl alcohol wash in a final concentration of 0.1 μg/μL to the tubes containing the isopropanol/NaCl mixture. After freezing at –30°C overnight, samples were centrifuged at 15,000 × g for 5 minutes, all liquid was removed by pipette, and pellets were left to air dry in a fume hood. Eluate was resuspended in 200 μL 1× TE Buffer, Low EDTA.

For all samples, the extracted eDNA yield was quantified (nanodrop) and samples were stored in 1.7 mL microtubes at –30°C until qPCR analysis. Biological replicated from the same location and depth were extracted and analyzed by qPCR separately before averaging. For details of all eDNA collections, see Table S3.

### qPCR of eDNA

To quantify the presence of neoplastic DNA and *M. arenaria* DNA in an eDNA sample, allele-specific qPCR primers targeting somatic SNVs in mitochondrial DNA were created. Based on previous genomic analysis of MarBTN (23), we identified loci on the mitochondrial DNA with two closely spaced SNVs that were found only in the USA sublineage of MarBTN, and were therefore somatic mutations that would be unique markers of cancer DNA. We designed qPCR primers (MarBTN-mt8807F1/MarBTN-mt8843R2) to specifically target these MarBTN-USA mutations (T8807G and C8843T), and we designed control primers (MarBTN-mt8807NORMF1/MarBTN-mt8843NORMR1) to amplify the same region, excluding the site of the SNVs. A single standard plasmid (pCR-MarBTNmt88/Norm) was cloned using the control primers to amplify the mitochondrial DNA of MarBTN cells (Zero Blunt Topo PCR cloning kit, Invitrogen). The plasmid was linearized using 0.5 μL of NotI-HF (New England Biolabs) for 2 h at 37°C in a 10 μL reaction, and heat-inactivated at 65°C for 20 min. DNA concentration was determined using Qubit, copy number per μL was calculated based on total plasmid size, and stocks were diluted to 1 × 109 copies/μL with NEB Elution Buffer. Standard curves were prepared through further dilution to tenfold reductions between 1 × 107 copies per reaction to 10 copies per reaction. For eDNA, 20 μL reactions were run with 2 μL of extracted DNA (using a higher volume of extracted DNA or a lower reaction volume led to errors due to PCR inhibitors in some samples). We used PowerUp SYBR green 2x master mix (ThermoFisher Scientific) with a StepOnePlus real-time PCR cycler (Applied Biosystems) with a holding stage at 95°C for 2 min, 40 cycles of 95°C for 15 s and 60°C for 30 s, followed by the melt curve stage. All samples were run in triplicate and averaged. Samples were considered positive if amplification was detected in all triplicate wells above 1 copy per reaction.

## RESULTS

### Identification and tracking of an outbreak of BTN in soft-shell clams in Triangle Cove and South Skagit Bay, Washington, USA

In 2022, when we sampled hemolymph from clams from Triangle Cove, Puget Sound, WA, we unexpectedly found that many clams (13 of 47; 28%) had abnormal cell morphology, consistent with DN (Figure 1). We confirmed that the DN in these clams was MarBTN using qPCR primers that amplify a characteristic retrotransposon insertion site in the clam genomes that is specific to this cancer lineage. We then tested a second location close to Triangle Cove (Stanwood, in southern Skagit Bay) and found cases of MarBTN there as well (5 of 39; 13%). We then sampled more broadly, collecting and diagnosing animals from 5 different beaches in Puget Sound, and found increasing prevalence of MarBTN at the two positive sites in 2023 and 2024 (Fig. 2, Table 1), with high severity of infection in many cases (Fig. S1). In the 2023 samples from Triangle Cove, we found 15 out of 50 animals (30%) to be triplicate positive by qPCR, and in 2024 we found 22 out of 27 animals (81%) to be positive (Table 1). In the 2023 samples from Stanwood, we found 11 out of 30 animals (36%) to be triplicate positive, and in 2024 we found 51 out of 66 animals (77%) to be positive (Table 1). The remaining 3 locations, Crandall Spit, Similk Bay, and Sequim Bay, did not have any triplicate positive amplification of cancer DNA from the animals sampled in 2023 or 2024 (Table 1).

**Figure 2.**
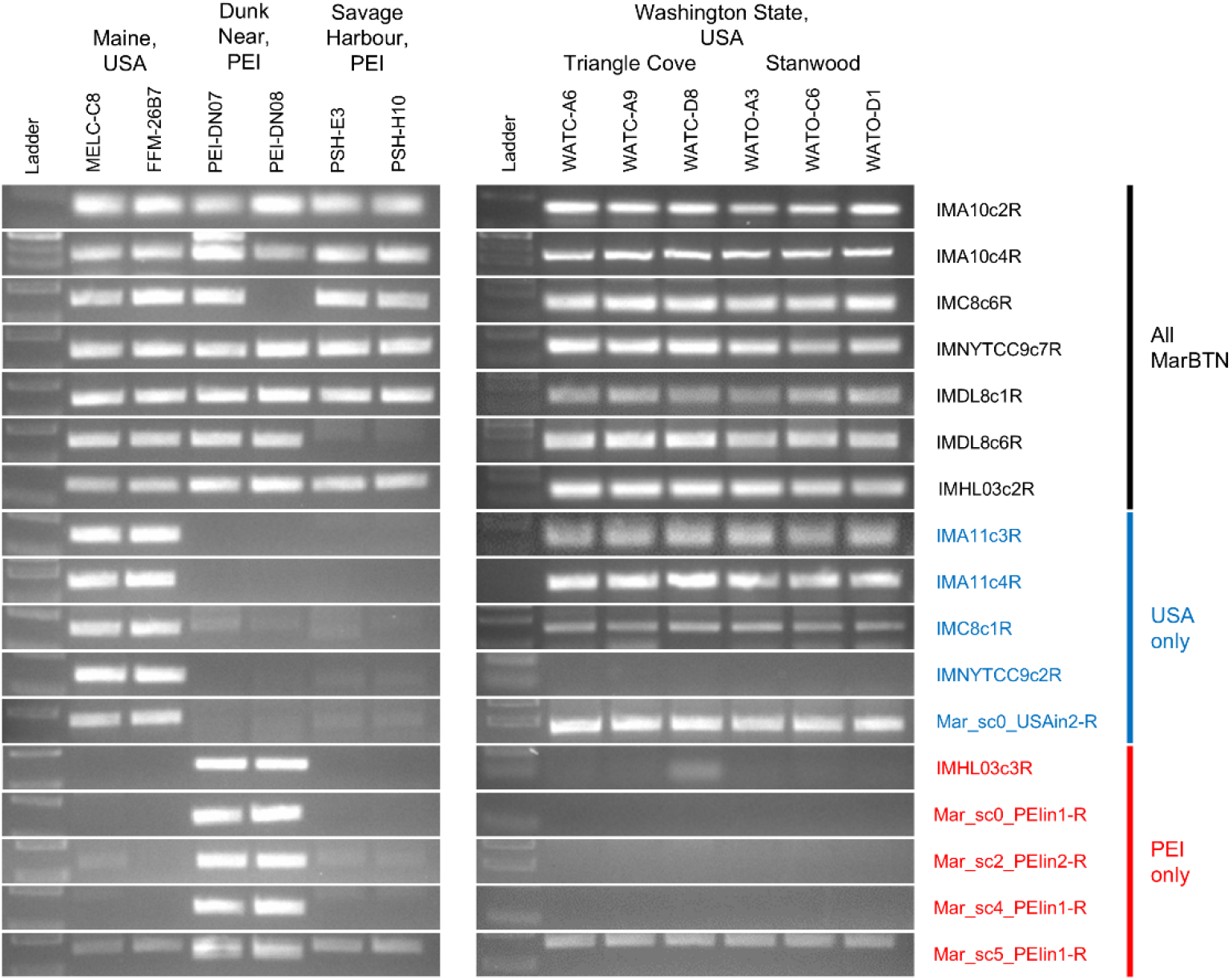
Transposon insertions show BTN in soft-shell clams in Puget Sound is from the USA sublineage of MarBTN. Multiple PCR primers diagnostic for specific insertion sites of the retrotransposon *Steamer* were used to amplify MarBTN DNA from clams in Puget Sound (three from Triangle Cove, WATC, and three from Stanwood, WATO) as well as from MarBTN DNA from clams known to carry the previously known USA sublineage (MELC and FFM), the previously known PEI sublineage (PEI), and new, more divergent samples collected from Savage Harbour, PEI (PSH). Labels on the right note the reverse primer used and whether the *Steamer* insertion sites have been previously observed in both USA and PEI sublineages (black), or were unique to USA (blue) or PEI (red) in previous studies (6, 23, 34).

**Table 1.** MarBTN prevalence in clams collected in Puget Sound, WA, USA, from 2022 to 2024.

| Location | Code | Coordinates | Collection date | Clams tested | MarBTN positive | Prevalence |
| --- | --- | --- | --- | --- | --- | --- |
| Triangle Cove | TC | 48.20013, -122.46748 | 4/21/2022 | 47 | 13 | 28% |
|  |  |  | 5/10/2023 | 50 | 15 | 30% |
|  |  |  | 7/8/2024 | 27 | 22 | 81% |
| Stanwood | ST | 48.24112, -122.37181 | 9/29/2022 | 39 | 5 | 13% |
|  |  |  | 8/2/2023 | 30 | 11 | 37% |
|  |  |  | 7/5/2024 | 66 | 51 | 77% |
| Crandall Spit | CS | 48.2941, -122.574528 | 5/6/2022 | 48 | 0 | 0% |
|  |  |  | 6/6/2023 | 34 | 0 | 0% |
|  |  |  | 6/20/2024 | 27 | 0 | 0% |
| Similk Bay | SM | 48.445834, -122.57675 | 7/6/2023 | 38 | 0 | 0% |
|  |  |  | 7/22/2024 | 15 | 0 | 0% |
| Sequim Bay | SQ | 48.0319, -123.01985 | 8/30/2023 | 3 | 0 | 0% |
|  |  |  | 2/7/2024 | 13 | 0 | 0% |

**Table 2.**
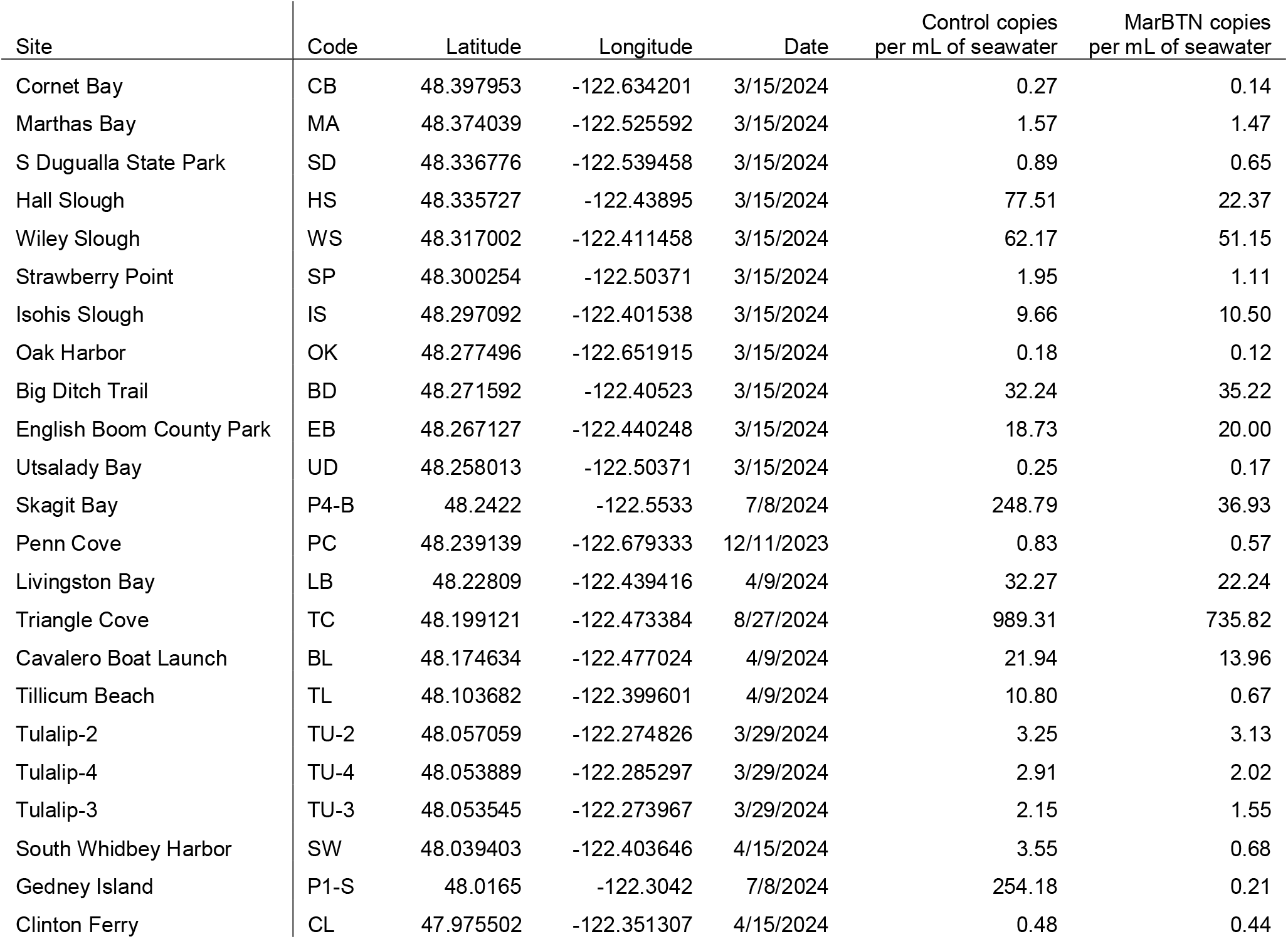
Sites in Puget Sound with positive detection of MarBTN in seawater using qPCR of eDNA.

### Molecular analysis suggests cancer in Puget Sound clams came from the East Coast of USA

The cancer detected in Puget Sound clams is likely from the same lineage as the MarBTN found in soft-shell clams on the east coast of the United States, based on detection of the cancer cells with the qPCR primers that amplify a *Steamer* transposon insertion specific to that lineage (Figure 2). In order to confirm this and to determine whether the cancer was transferred to the west coast from one of the existing sublineages of MarBTN from USA or PEI or whether it diverged prior to the split of those two sublineages, we used 17 primers that are diagnostic to specific *Steamer* integration sites. Of these primers, seven were previously identified to be present in MarBTN samples from both PEI and USA, five were specific to samples from USA, and five were specific to samples from PEI (see Supp. Table 1). As controls, we tested two samples of MarBTN from the east coast of the USA and four from PEI clams (two collected previously (6) and two recently collected). We then analyzed six samples of MarBTN from the west coast (three each from Triangle Cove and Stanwood). All of the DNA samples from control samples and west coast MarBTN amplified with the primers that target insertions previously known to have occurred prior to the USA/PEI split, with the exception of PSH-H3 and PSH-E10 with primer IMDL8c6R (Fig. 3), which is likely due to a deletion of that region. The PEI specific primers amplified in the two samples previously collected from the Dunk River Estuary, PEI (PEI-DN07 and PEI-DN08), but did not amplify in west coast samples or in the two newer samples from Savage Harbour, PEI (PSH-H3 or PSH-E10), suggesting that these newer PEI samples may reflect a new sublineage that branched off from the previously known sublineage in PEI. The USA-specific primers amplified all of the east and west coast samples except for IMNYTCC9c2R, which did not amplify in the west coast samples (Fig. 3). We additionally amplified and sequenced the mitochondrial cytochrome b oxidase I (*mtCOI*) locus from these samples, and we confirm that the sequence is identical to that previously found in the USA sublineage of MarBTN, without the SNV specific to PEI MarBTN. Therefore, the MarBTN in Puget Sound, on the west coast of the USA closely matches known samples from the USA sublineage, and likely was recently transported from New England to the location of this new outbreak.

**Figure 3.**
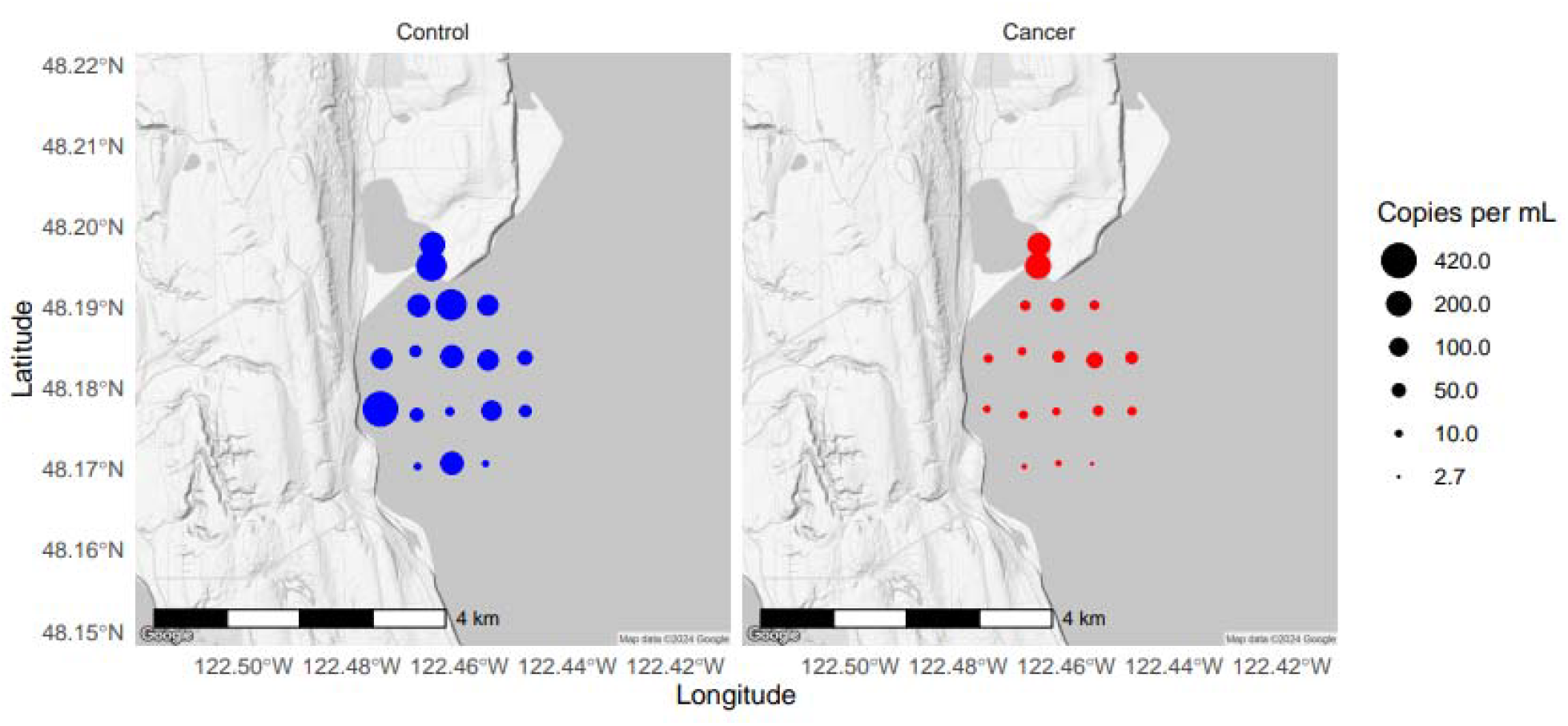
Detection of BTN in environmental DNA (eDNA) at a site of disease outbreak (Triangle Cove, Camano Island, WA) Samples of seawater were taken by boat at the entrance to Triangle Cove, Camano Island, Washington, and at multiple sites further from the entrance. The amplification from control primers (which amplify all *Mya arenaria* mitochondrial DNA, left) and from cancer-specific primers (which target two somatic mutations unique to the USA-sublineage of MarBTN, right) are shown, with sizes shown by the size of the point. Points are the average of two samples taken in a single day. Absolute quantification was calculated as copies per reaction using a standard plasmid control, normalized to copies per mL of seawater, assuming 100% efficiency of eDNA extraction from water samples.

### Sensitive mitochondrial qPCR on eDNA from Triangle Cove, Washington, shows local spread of cancer cells in seawater

Collection and diagnosis of clams is highly sensitive, but requires considerable work at each collection that limits the widespread use of this method to track the spread of transmissible cancer across large areas. Environmental DNA (eDNA) methods have been used to detect MarBTN-specific DNA in aquarium water of diseased clams, but the sensitivity only allowed limited detection in wild samples (17). We therefore developed specific qPCR primers that target SNVs in the mitochondrial genome of MarBTN that were previously found only in the USA sublineage (23). These SNVs are somatic mutations, and they would be unlikely to be found in any healthy host mitogenomes. As there are many copies of the mitochondrial genome per cell, these targets could be much more sensitive than primers targeting nuclear loci. They were first tested using water collected from the site of the outbreak at Triangle Cove, WA. Water samples at the entrance to the cove were strongly positive, with lower levels of MarBTN detected in the seawater at greater distances from the entrance to the cove (Figure 3). Interestingly, the control primers, which amplify the same mitochondrial locus targeting conserved sites, amplified to higher levels than the cancer primers at some of the sites furthest from the cove, suggesting that either healthy clam cells can survive in the environment longer than MarBTN or that there are populations of clams with lower prevalence outside of the cove itself. These data show that transmissible cancer cells can be detected through eDNA, and that while the amount of MarBTN decreases over distance, it can be detected as far as 2 km from the putative source of the cells.

### Surveys using sensitive mitochondrial qPCR on eDNA shows the spread of MarBTN throughout Puget Sound, Washington, USA

Extraction of eDNA from seawater samples, followed by qPCR targeting the mitochondrial somatic SNVs was used to detect MarBTN in environmental sites throughout Puget Sound to determine the spread of MarBTN throughout the region, without the need for identifying specific clam populations or collecting and diagnosing individual animals. Water samples were collected from shore at five sites in 2023 and water samples were collected by boat as well as from shore from 47 sites from 15 Mar through 27 Aug, 2024 (Figure 4, Supplementary Table 3). Amplification with the control primers identified soft-shell clam DNA at most water collection sites except for those farthest from shore and at two locations along the west coast of Whidbey Island (Figure 4A). The highest levels of control amplification could be found at the known sites where we had previously collected clams, as well as some additional areas, such as southern Whidbey Island, around Everett, and further north in Skagit Bay, suggesting significant soft-shell clam populations in those areas. As expected, MarBTN was detected at high levels at Triangle Cove and Stanwood, but it was also observed at multiple sites further north into Skagit Bay, further south in Port Susan, and southwest Whidbey Island (Figure 4A,B). These new data reveal an outbreak beyond the initial sites previously observed, but also show that currently only a small part of the total soft-shell population in Puget Sound is affected, and several large populations that are not yet impacted by MarBTN exist on either side of the current outbreak.

**Figure 4.**
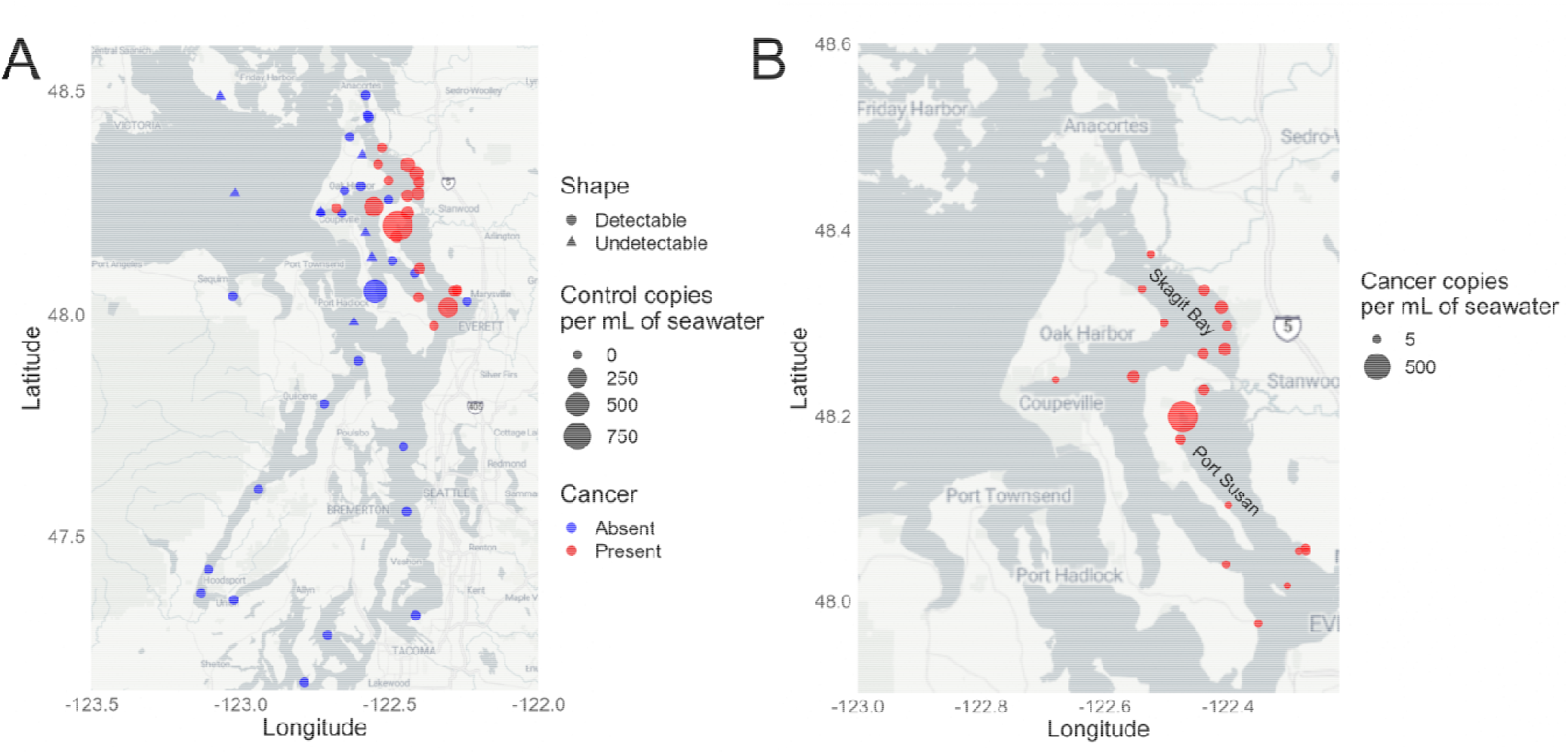
Spatial distribution of MarBTN outbreak shown by broad eDNA surveys across the Puget Sound. In 2023 and 2024, samples of water were collected from 50 locations in Puget Sound, Washington. Environmental DNA (eDNA) was extracted and samples were tested for the presence (circle) or absence (triangle) of DNA from soft-shell clams (*Mya arenaria*), using control primers. (***A***) The points on the map show collection locations and the size of the point shows the magnitude of the control amplification. The color denotes whether cancer-specific DNA was detected (red) or absent (blue). (***B***) A more detailed map of the area of the outbreak shows all points where BTN was detected, and the size of the points show the quantity of the MarBTN-specific amplification. Absolute quantification was calculated as copies per reaction using a standard plasmid control, normalized to copies per mL of seawater, assuming 100% efficiency of eDNA extraction from water samples. Maps were generated using R packages ggplot, ggmap, and dplyr. Map style was generated by StadiaMaps, using map type “alidade_smooth.”

## DISCUSSION

Through the analysis of clam hemolymph and a novel sensitive eDNA assay, we report the identification of an outbreak of MarBTN in soft-shell clams on the west coast of the United States, a population previously believed to be unexposed to the disease. Multiple genetic markers confirm that the MarBTN in west coast clams is most closely related to the USA sublineage, which was previously thought to be found only on the east coast of the US. The lack of previous detection, rapidly increasing prevalence, and small geographic area affected all strongly suggest that this disease was recently transplanted from the east coast of the United States to the west coast, initiating this outbreak. The long-distance transfer of MarBTN is likely due to an accidental transplantation of an infected clam or accidental transfer of seawater containing cancer cells. This would be the second case of a BTN spread to new oceans by human activities after the MtrBTN2 lineage that has been observed in mussels in the *Mytilus* genus worldwide (8, 11).

There have been significant outbreaks of disseminated neoplasia on the east coast of the United States, of up to 90% prevalence (21, 22) that correlated with severe population die-offs and are likely to have been caused by MarBTN. However, this disease is currently maintained in clam populations on the east coast at low enzootic levels throughout all known populations in the region and has not led to any recent severe population losses. Moreover, based on genomic analysis of MarBTN, this lineage was estimated to be over 200 years old (23), showing that it has been evolving together with east coast clams for many generations. This newly reported outbreak of MarBTN in west coast clams would be expected to follow a similar pattern in the future as the outbreak progresses, with significant mortality and severe population loss, followed by selection for more resistant clams, and finally enzootic maintenance of the disease in the population. The potential for local populations to face extirpation from this disease remains unclear, as does the timeline or nature of any potential evolutionary resistance.

This study is the first of its kind to use eDNA sampling to detect transmissible cancer in wild seawater samples. The mitochondrial-targeted eDNA method enabled sensitive tracking of the outbreak over a large geographic area. It will be critical to following disease spread in the future, and it could be applied to other BTN lineages in other species. The rate at which BTNs spread from one population to another along a coast is currently unknown. BTNs have only recently been identified as an infectious disease, and this eDNA method will be vital for observing disease dynamics in the environment over time, and the recognition of this outbreak at an early stage make it a critical case for understanding the mechanisms and speed of spread of this type of infection in the environment.

## ACKNOWLEDGEMENTS

We thank Megan Dethier (Friday Harbor Laboratories, University of Washington) for advice and help in locating *Mya arenaria* in Puget Sound and the Salish Sea, and Drew Harvell (Cornell University) for help in collecting the seawater sample from False Bay, San Juan Island. Ryan Kelly provided modified CTAB protocols. Funding comes from an NSF Ecology and Evolution of Infectious Disease grant (2208081) to MJM, JD, and an REU student (DL); an NIH training grant T32-HG000035 to SFMH; and NSF grant OCE-2349136 to fund the REU students at WWU (AS and HM).

## SUPPLEMENTARY DATA

Figure S1. Quantification of disease severity in soft-shell clams with MarBTN

Table S1. Soft-shell clam collection and diagnosis for MarBTN in multiple sites in Puget Sound, WA, USA

Table S2. Primers used

Table S3. Quantification of MarBTN in eDNA from seawater collected from multiple sites in Puget Sound, WA, USA

