## Supplementary material for "Identification of an Outbreak of Bivalve Transmissible Neoplasia in Soft-Shell Clams (*Mya arenaria*) in the Puget Sound Using Hemolymph and eDNA Surveys": Figure S1

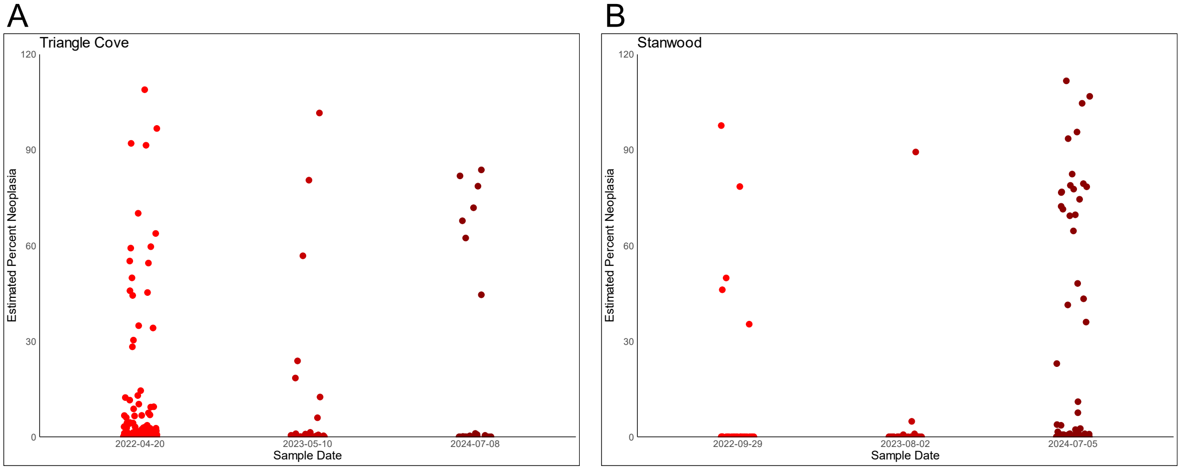


**Figure S1. Quantification of disease severity in soft-shell clams with MarBTN**

Soft-shell clams (*Mya arenaria*) from two sites in Washington State were found to be positive for MarBTN based on collections from 2022-2024: (***A***) Triangle cove, and (***B***) Stanwood (locations marked in map in Figure 1 and coordinates in Table 1). MarBTN diagnosis was made by qPCR analysis of hemolymph from collected clams, using a primer pair specific to a nuclear marker in MarBTN and a pair that amplifies that locus from all known soft-shell clam alleles. The estimated percent neoplasia in the hemolymph is shown, with a dot for each clam sampled.
